# Determination of best Raman spectroscopy spatial offsets for transcutaneous bone quality assessments in human hands

**DOI:** 10.1101/2021.07.26.453893

**Authors:** Keren Chen, Christine Massie, Hani A. Awad, Andrew J. Berger

## Abstract

Spatially offset Raman spectroscopy (SORS) is able to detect bone signal transcutaneously and could assist in predicting bone fracture risk. Criteria for optimal source-detector offsets for transcutaneous human measurements, however, are not well-established. Although larger offsets yield a higher percentage of bone signal, the absolute amount of bone signal decreases. Spectral unmixing into bone, adipose, and non-adipose components was employed to quantify changes in bone signal to noise ratio across a range of offsets, and optimal offsets for phalanx and metacarpal measurements were determined. The bone signal to noise ratio was maximized at offsets ranging from 4-6 mm.

## Introduction

Osteoporosis is a worldwide concern that significantly affects post-menopausal women and the aging population of both sexes [1]. This bone disease is characterized by reduced bone mass, deterioration of the bone’s micro-architecture, and decreased bone strength [2]. The reduction in bone strength will increase a person’s risk of experiencing a fragility fracture. After suffering an osteoporotic fracture, people are then likely to experience morbidity, other fragility fracture, and even, sometimes, mortality [3]. Bone mineral density (BMD) is measured through dual-energy x-ray absorptiometry (DXA) to diagnose osteoporosis. However, many at risk patients are not screened by DXA before experiencing a fracture due to health care disparities and accessibility issues [4]. Furthermore, DXA does not accurately identify patients who are at high risk of experiencing a fragility fracture [5], largely because it senses only bone mineral, providing no information about the relative abundance of bone matrix. Methods that are sensitive to a bone’s biochemical composition may aid in fracture risk screening and prediction, since the composition is related to the bone’s biomechanical properties [6-8].

Raman spectroscopy is a non-invasive, vibrational technique that can measure the biochemical signature of bone. The ratio of mineral and matrix Raman signal strengths is a common metric to characterize bone health. However, limited progress has been made in using Raman spectroscopy for transcutaneous *in vivo* bone quality assessment due to the unavoidable spectral contribution from the surrounding soft tissues. Any transcutaneous Raman measurement will contain type I collagen Raman signals from both bone and soft tissue, making it difficult to determine the true chemical balance of mineral and matrix in the bone alone. Spatially offset Raman spectroscopy (SORS) transcutaneous measurements can increase specificity to bone’s spectral features *in vivo*, but the contribution from overlying tissue is not eliminated.

Bone quality metrics derived from Raman spectroscopy have been explored in rodent models both *in vivo* and *ex vivo* on tibiae and femurs [9-13]. In these models the bone is typically 1 mm from the tissue surface. The soft tissue overlying human tibiae and femurs is thicker than the soft tissue in rodents, however, thus requiring deeper optical penetration to acquire sufficient bone signal. The thickness of soft tissue overlying human tibiae is 4.8 ± 2.0 mm [14]. Human fingers have significantly less overlying soft tissue (2.0 ± 0.7 mm), making them a better candidate location for initial *in vivo* SORS measurements. In addition, from a clinical standpoint, the hand is already an explored candidate for clinical bone quality assessment. Specifically, radiographs of metacarpals have been demonstrated to predict vertebral fracture risk in post-menopausal women [15], predict hip fracture risk [16], and correlate to hip bone mineral density (BMD) [17].

Although SORS can detect signals from phalanges *in vivo* using various offsets from 0-10 mm [18, 19], it remains unclear what metric should guide the choice of SORS offsets. Larger offsets always increase the ratio of the signal from the bottom layer to that of the top layer, but as we have previously noted, the signal-to-noise ratio (SNR) for the bottom layer will eventually decrease when the offset is too large [20]. For a given specimen, there is thus an offset at which bone SNR reaches a maximum. This distance will be affected by the thickness of the overlying soft tissue, as this is the dominant source of fluorescence and noise in transcutaneous SORS measurements. Thickness will depend upon the type of region explored. While both metacarpals and phalanges may be suitable for bone quality assessment, the soft tissue overlying metacarpals is typically thicker than phalanges. Variation in body mass index (BMI) values between subjects will also cause differences in soft tissue thicknesses overlying the same type of bone. These factors will influence the choice of source-detector offsets for SORS measurements. Here, we investigate how the ratio of bone signal to noise varies for (a) different regions within human hand cadavers and (b) between hands with different BMI values in order to determine an optimal source-detector separation range.

In addition, we consider the need for a more sophisticated spectral unmixing model. In our previous work on murine tibiae, the Raman spectral lineshape of soft tissue was treated as spatially invariant within the optically interrogated SORS volumes. Due to the greater thickness of the overlying soft tissue and the larger SORS offsets required in this study, measurements were more sensitive to spatial heterogeneity in the soft tissue. We therefore employed a three-tissue model in this work, allowing for two different sub-types of soft tissue (adipose and non-adipose) along with bone.

## Methods

### 1.1 Cadaver samples

We obtained two cadaver hand samples from the Anatomy Gifts Registry (Hanover, MD). One sample came from a Caucasian 61-year-old male with a healthy BMI of 24. The other sample was from a Caucasian severely obese 61-year-old female with a BMI of 36.

### 1.2 Raman spectroscopy

All the Raman spectra in this study were acquired using an 830-nm, 120-mW laser focused in a spot with a diameter of about 1.0 mm for 300 s. Measurements were performed transcutaneously over metacarpals II and III (henceforth M2 and M3, respectively) and phalanges II and II (P2 and P3), as shown in Fig.1(a). Fig. 1(b) sketches the main optical components and demonstrates how source-detector offset was varied by linearly translating the excitation delivery arm. At each of the finger locations SORS measurements were taken at offsets ranging from 0-8mm. Spectral pre-processing consisted of cosmic ray removal, CCD readout and dark current subtraction, and image aberration correction [21]. The spectra were fit to a 5^th^ order polynomial to remove fluorescence and smoothed with a Savitzky-Golay filter [22] over a window of 5 cm^-1^. Spectra were normalized to their mean absolute deviation (MAD) [13].

**Fig.1.**
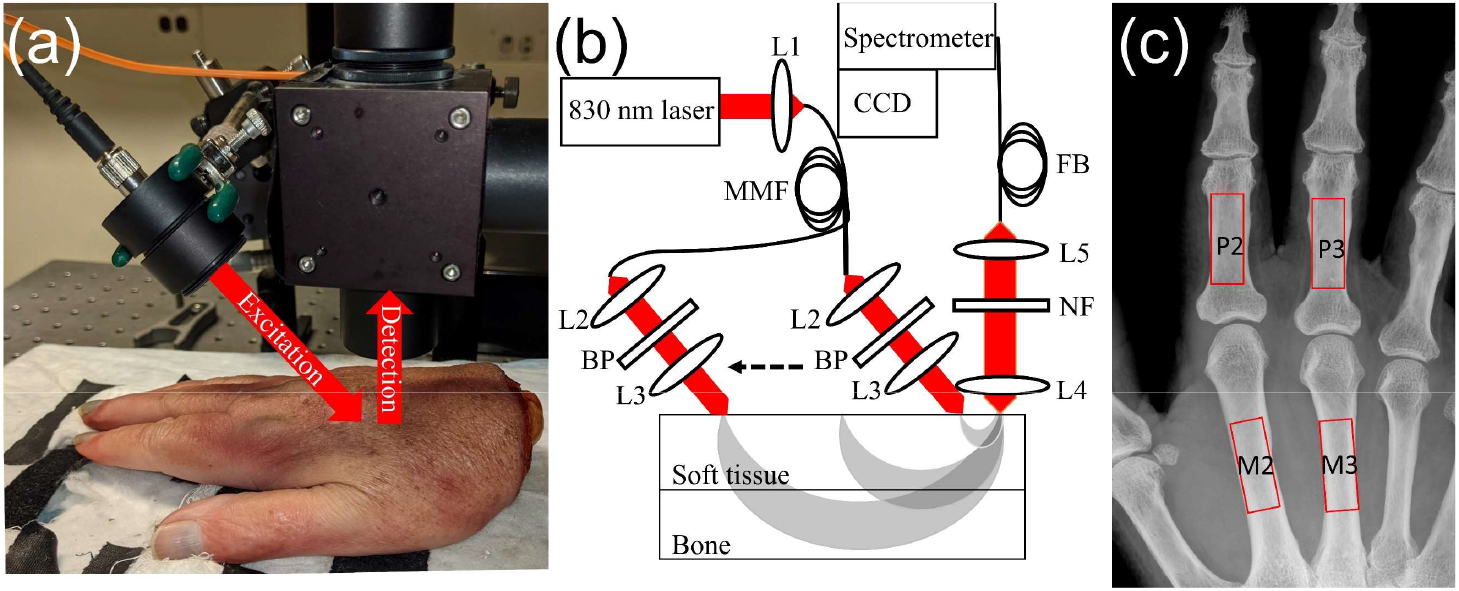
(a) Setup for human cadaver measurement with separate laser-focusing and Raman-collecting optics. (b) Schematic of the optical system. Shaded regions indicate how the mean penetration depth changes as the source-detector offset is varied by linear translation of the laser illumination spot. L1, L2, L3, L4 are lenses. BP is a band-pass filter for 830 nm. NF is a notch filter blocking 830 nm. (c) X-ray image of one hand specimen. Anatomical measurement positions are marked with red boxes.

Figure 1(a) shows the setup of the system used in this study. The excitation optics focusing the laser beam on the skin surface are mounted on a translation stage that moves the laser spot parallel to the finger’s orientation. While moving the excitation probe away from the collection end, i.e. with larger offset, a higher fraction of the collected Raman scattering originates from the deeper layer. The schematic of the system and SORS measurement geometry are shown in Fig. 1(b). Fig. 1(c) shows a X-ray image acquired from one hand specimen. The interrogating locations P2, M2, P3 and M3 are marked with red boxes on the figure.

### 1.3 Data analysis

To quantitatively analyze the spectral composition, a simultaneous, overconstrained, library-based decomposition (SOLD) method we introduced before [23, 24] was applied to quantify the separate spectral contributions from bone, fat, and non-adipose soft tissue regions. The SOLD algorithm drew upon spectral libraries (one for each of the three tissue categories) to estimate basis spectra that best fit the multiple-offset SORS data for a particular finger region; each location was analyzed separately. The previously-acquired libraries consisted of 102 bone spectra and 159 soft-tissue spectra measured from murine tissue specimens. The soft-tissue spectra were subsequently separated into adipose and non-adipose libraries. The SOLD process produced (a) sample-specific basis spectra of bone, adipose, and non-adipose soft tissue for each region and (b) corresponding weight coefficients for the three tissue types at each source-detector offset. Any single transcutaneous spectrum, at any offset, could thus be unmixed into a weighted superposition of three basis spectra. Areas under individual Raman peaks could be tracked as a function of spatial offset.

Signal and noise levels were calculated as follows. Spectral intensity counts were converted to photoelectron numbers by multiplying by the CCD conversion factor (e-/counts, 2.5 for this detector array). Raman peak areas were calculated from 932 cm^-1^ to 992 cm^-1^ (phosphate), 1384 cm^-1^ to 1504 cm^-1^ (CH_2_) and 1626 cm^-1^ to 1686 cm^-1^ (Amide I); these quantities defined “signal”. The square root of the total counts (Raman plus broadband emission from fluorescence and other effects) in each of these spectral regions was defined as the corresponding “noise”. By this process the ratios of bone-to-noise (BNR), adipose-to-noise (ANR), and non-adipose-to-noise (NANR) were calculated using the areas under the phosphate peak of bone, the CH_2_ peak of adipose, and the Amide I peak of non-adipose, using the unmixed spectra estimated by SOLD for each source-detector offset separately.

To investigate the number of independent spectral components needed to model the SORS data, principal component analysis (PCA) was performed upon the nine spectra gathered from the P2 region of the low BMI specimen. This produced orthonormal loading spectra that could be examined for Raman-like features above the shot noise.

## Results

Raman spectra measured from the P2 region of the cadaver with healthy BMI, with source-detector offsets from 0 to 8 mm, are shown in Fig. 2 (a). All spectra exhibit prominent soft tissue and bone peaks such as Amide I (1656 cm^-1^), CH_2_ (1450 cm^-1^), Amide III (1243-1320 cm^-1^), hydroxyproline (876 cm^-1^), carbonate (1070 cm^-1^) and phosphate (959 cm^-1^). As the offset increases the absolute signal level goes down, but as expected the percent contribution from subsurface bone mineral increases. This effect is more evident in Fig. 2(b), which replots the spectra between 4 and 8 mm with the area of the phosphate peak region normalized (dashed box, 932 cm^-1^ to 992 cm^-1^). In this rendering the Raman signals from the amide and CH_2_ regions noticeably decline versus offset. The shot noise amplitude in the spectrally flat region near 1200 cm^-1^ (solid box), however, noticeably *increases*, making the bone signal to noise ratio worse. Optimizing the offset for best phosphate signal to noise is further complicated by the overlapping presence of peaks from organic components (shoulder at 940 cm^-1^).

**Fig.2.**
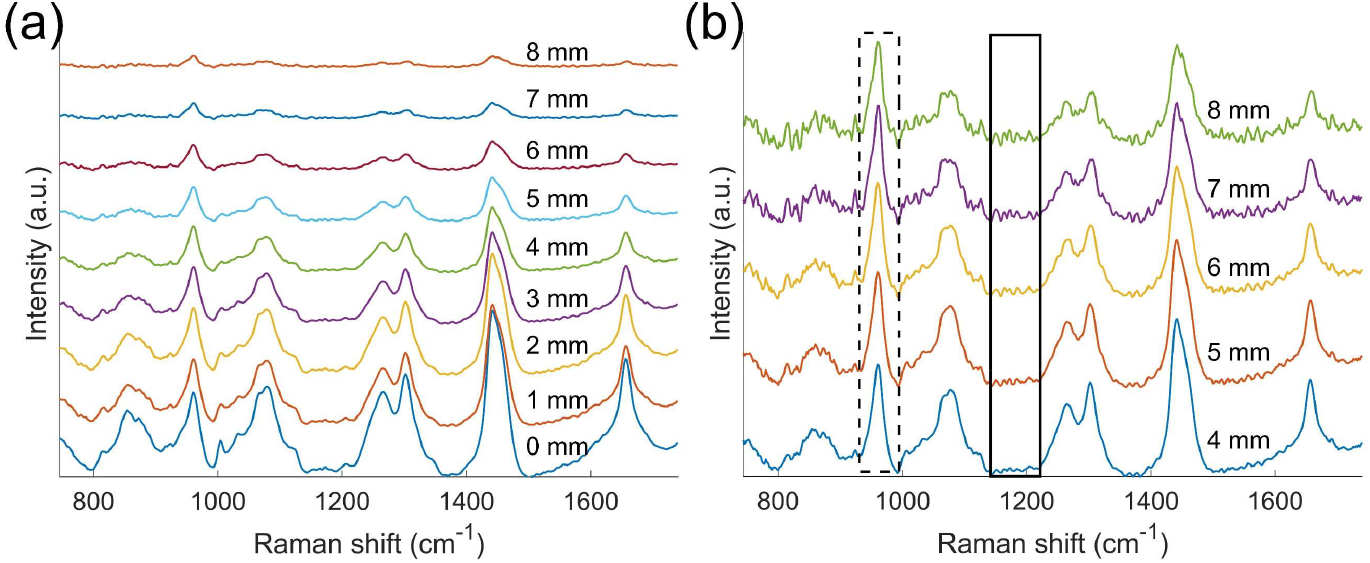
(a) Raman spectra acquired with offset from 0 mm to 8 mm on P2 from the healthy-BMI cadaver specimen. (b) Raman spectra normalized to area under the curve in the phosphate region (marked with dashed rectangle), with noise region marked with solid rectangle.

As noted earlier, the transcutaneous spectra were unmixed to isolate spectral contributions from each tissue type. Fig. 3 shows representative unmixing results from nine offset measurements (0 to 8 mm) from the M2 region of the cadaver with healthy BMI. SOLD analysis generated basis spectra for bone (c), non-adipose soft tissue (d), and adipose tissue (e) that best described the nine transcutaneous Raman spectra. As an example of the fitting process, the 3-mm offset spectrum is shown (a) along with the best basis-spectrum model (b) and the corresponding residual (f).

**Fig.3.**
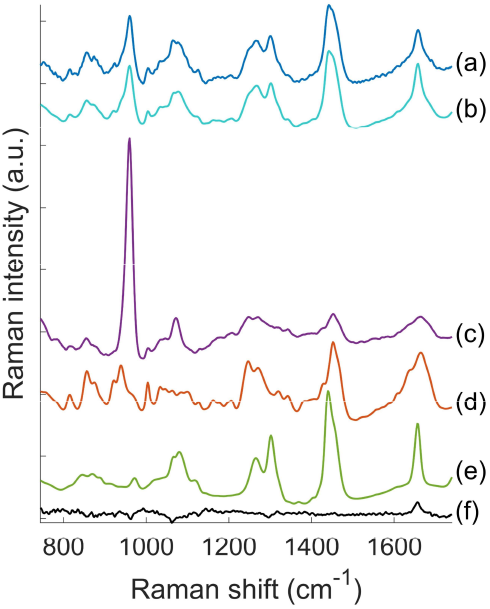
Three layer SOLD processing. (A) 3 mm offset transcutaneous measurement from M2 region on the healthy-BMI cadaver. (B) best spectral fit to A using SOLD. SOLD-estimated spectral contributions in B from bone (C), non-adipose (D), and adipose tissues (E), respectively. (F) residual after SOLD fitting, which is A subtracted by B.

Figure 4(a) plots the calculated phosphate to Amide I ratio (PTAR), phosphate peak area under curve (AUC) from unnormalized data, and phosphate to noise ratio (PNR) as a function of source-detector offset for a P2 region of the cadaver with healthy BMI. Differing trends are seen for PTAR and phosphate AUC. The PTAR, which reports the relative abundances of phosphate and collagen in the interrogated tissue region, increases monotonically with offset. The phosphate AUC, which reports the absolute (unnormalized) strength of phosphate signal, shows the opposite trend, *decreasing* monotonically with offset. Due to these competing effects, PNR—the signal to noise ratio of the bone’s spectral contribution---maximizes in the middle of the explored range of offsets, around 3-5 mm.

**Fig.4.**
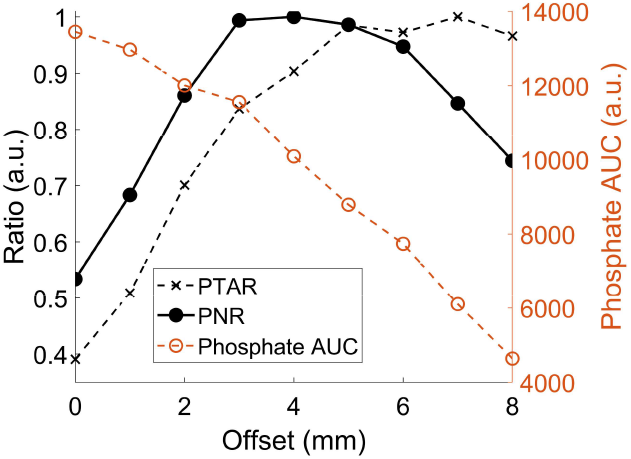
Phosphate to Amide I ratio (PTAR), phosphate to noise ratio (PNR) and phosphate peak area under curve (AUC) of P2 versus offset from the healthy BMI specimen, normalized to their maximum values.

Figure 5(a) compares the SNR trends vs. offset for all three tissue types for the P2 region on the healthy BMI specimen, with dashed vertical lines marking out the average of maxima for each ratio. Non-adipose soft tissue to noise ratio (NANR) decreases monotonically with offset, while adipose to noise ratio plateaus at 2—3 mm offset. Figure 5(b) plots the trends for the four different regions on the healthy BMI specimen. Soft tissue trends are visually comparable for all regions. For non-adipose tissue, the average offset of the four maximum NANR values is ∼0.5 mm; for adipose, it increases to 2mm. The BNR trends were less comparable between the phalanges and metacarpals and were therefore averaged separately. The average maximum-BNR offset for the two phalanges was 4 mm; for metacarpals, it increased to 5.5 mm.

**Fig.5.**
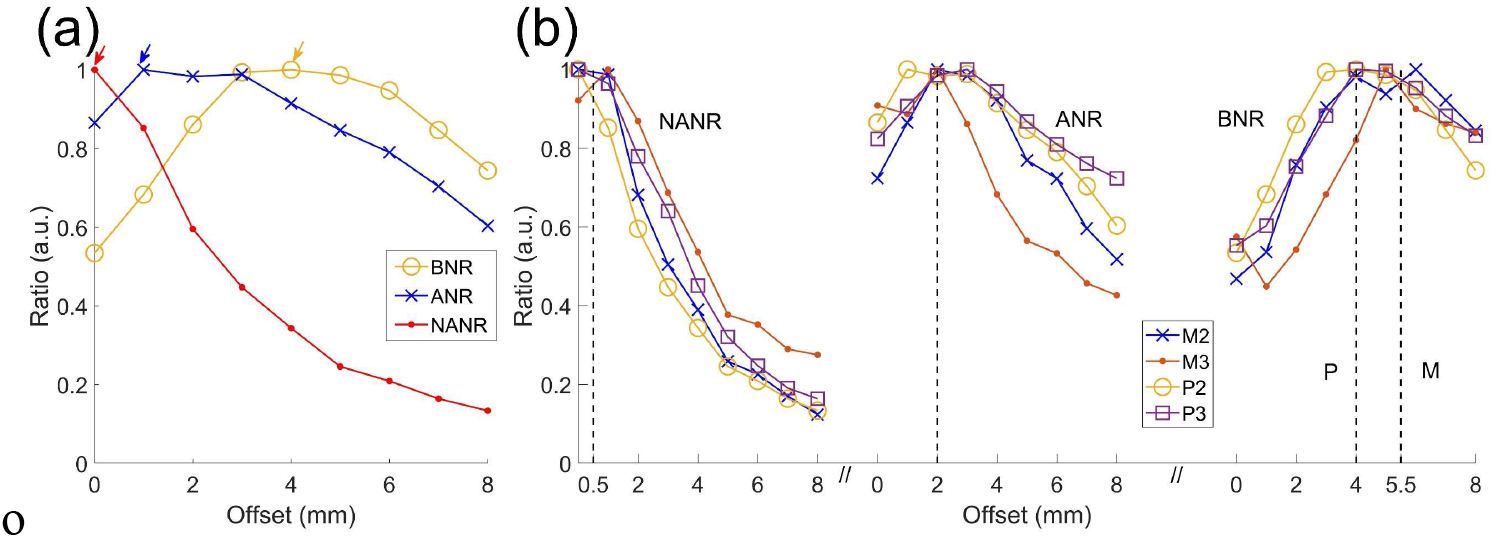
Quantification of data collected from healthy BMI subject. (a) comparison between trends of non-adipose to noise ratio (NANR), adipose to noise ratio (ANR) and bone to noise ratio (BNR) and acquired from P2 region. Maximum ratios are marked with arrows. (b) NANR, ANR and BNR from multiple locations (M2, M3, P2 and P3). Averaged maxima are marked with dashed lines.

BNR values for two hand specimens with different BMI levels [24 (healthy) and 36 (severely obese)] are compared in Fig. 6. For both the P2 and M2 regions, the lower BMI specimen produced the larger BNR value. For each specimen, the BNR values was larger for the phalanx than the metacarpal.

**Fig.6.**
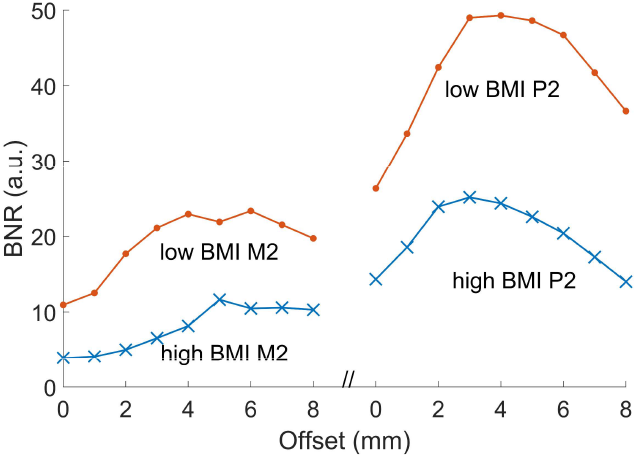
Comparison of bone to noise ratio acquired from M2 and P2 of two hands with different BMI (24 versus 36).

Figure 7 shows the loadings of the first four principal components (PCs) from PCA of the spectra from the P2 region of the low BMI specimen. The loadings of PC1 (which explains 84.99% of the spectral variance) clearly exhibit the phosphate peak 959 cm^-1^. PC2 (11.30%) and PC3 (2.22%), while more complex, exhibit upward and downward Raman-like features above the noise, particularly in PC3. PC4 (0.50% variance), by comparison, is visually dominated by noise.

**Fig.7.**
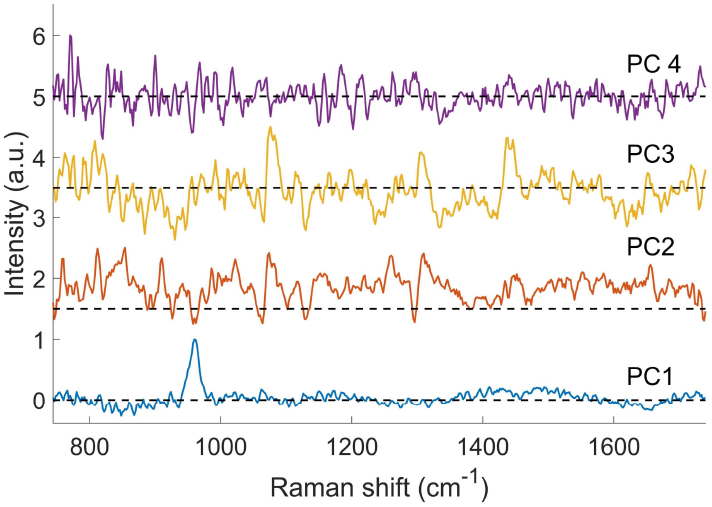
Loadings of PC1, PC2, PC3 and PC4 normalized to their maximum values, derived from offset measurements acquired from P2 of low BMI hand, using PCA.

## Discussion

This study investigated the use of SORS for eventual *in vivo* assessment of bone quality in human hand bones. In transitioning from mouse measurements to larger size scales needed in human measurements, the sophistication of the spectral unmixing model and the selection of source-detector offsets needed to be reassessed.

The spectral data suggested the need for a three-tissue unmixing model when measuring human metacarpals and phalanges *in situ*. Specifically, Raman scattering from non-adipose and adipose soft tissues exhibited different trends in SNR as the source-detector offset was increased (Figure 5). Non-adipose SNR attained its largest values near zero offset, while adipose SNR values were largest with a few mm of separation. These results match expectations based upon anatomical considerations. Since non-adipose soft tissue constitutes 100% of the cutaneous tissue, adipose tissue has a deeper mean subcutaneous location. A zero-offset measurement is thus highly specific to non-adipose soft tissue, while a few-mm offset provides a slightly deeper mean sampling depth that is better for sensing the adipose distribution. We note that the trend lines shown in Figure 5 were derived independently, without any anatomical assumptions.

The different trends for adipose and non-adipose tissue suggest that a spectral unmixing model for human hands requires a separate basis spectrum for each tissue type. This is different from the transcutaneous murine model we have used previously, which assumed spatial invariance of the soft tissue spectral lineshape. The greater thickness of the soft tissue overlying human hand bones makes this assumption of soft tissue homogeneity invalid. Separately, the PCA results (Fig. 6) further suggest three independently varying basis spectra above the noise floor.

Over source-detector offsets ranging from 0 to 8 mm, our measurements exhibited maximal BNRs in the 3-5 mm range, over a variety of measurement locations (metacarpals and phalanges) on two cadaver hands with different BMI. As Fig. 4 emphasizes, this local maximum in BNR arises from two opposing trends. As source-detector separation gets larger, the absolute number of detected phosphate Raman photons *decreases*, but the fraction of total detected photons in that wavenumber bin (both Raman and fluorescence) that arises from phosphate Raman scattering *increases*.

In addition, a second hand with thicker overlying tissue (higher BMI) required larger offsets to get the best bone to noise ratio, both for metacarpals and for phalanges (Fig. 6). This matches the general understanding that SORS measurements with larger offsets provide a larger mean interrogation depth. In a similar comparison based upon thickness, it is reasonable that each cadaver’s phalanx BNR value was larger than for the metacarpal, since phalanges will tend to have the thinner soft-tissue layer.

Meanwhile, it should be noted that the libraries we used to decompose the transcutaneous spectra were acquired from tissues excised from mice in our previous study. Although visually the Raman spectra of human and murine tissues strongly resemble each other, a dedicated human tissue library could improve the performance of the fitting and further facilitate the analysis of spectra, which may help determining the best offset.

The maximum permissible exposure on living human tissue is more restricted than in this study. Here, by using an 830 nm laser with 120 mW the irradiance was 1.25 W/cm^2^, which exceeded the ANSI maximum permissible exposure of 0.363 W/cm^2^. In future studies, to compensate for the lower irradiance requirement, the exposure time could be increased and/or irradiance could be delivered to two separate regions of the same bone simultaneously.

In conclusion, this work addresses issues for transcutaneous Raman spectroscopy of human bones. The hand was selected as a target location, based upon the bones’ proximity to the skin surface and the accessibility of the hand for possible *in vivo* clinical applications. In switching from murine to human specimens, three spectral libraries rather than two were required to model the Raman peak variations in the SORS measurements; this was due to the larger source-detector offsets inducing more sensitivity to tissue heterogeneity. BNR, the ratio of phosphate Raman signal to noise, was introduced as a metric to optimize collection of Raman scattering from bone under a given set of measurement conditions. Optimizing BNR should be valuable in building Raman models to predict gold-standard bone metrics currently provided by X-ray and biomechanical tests.

## Conflict of Interest Statement

The Authors have no conflicts to disclose.

## Acknowledgements

We would like to thank Ms. Emma Knapp for assistance in obtaining hand specimens. The study was supported by grant numbers R01AR070613, R21AR061285, and P30AR069655 from NIAMS/NIH. The content is solely the responsibility of the authors and does not necessarily represent the official views of the NIH.

## Notes

### Competing Interest Statement

The authors have declared no competing interest.

## References

1. Riggs, B.L. and L.J. Melton, 3rd, The worldwide problem of osteoporosis: insights afforded by epidemiology. Bone, 1995. 17(5 Suppl): p. 505s–511s.

2. Coughlan, T. and F. Dockery, Osteoporosis and fracture risk in older people. Clinical medicine (London, England), 2014. 14(2): p. 187–191.

3. Nazrun, A.S., et al., A systematic review of the outcomes of osteoporotic fracture patients after hospital discharge: morbidity, subsequent fractures, and mortality. Ther Clin Risk Manag, 2014. 10: p. 937–48.

4. Hamrick, I., et al., Health care disparities in postmenopausal women referred for DXA screening. Fam Med, 2006. 38(4): p. 265–9.

5. Choksi, P., K.J. Jepsen, and G.A. Clines, The challenges of diagnosing osteoporosis and the limitations of currently available tools. Clinical diabetes and endocrinology, 2018. 4: p. 12–12.

6. Burstein, A.H., et al., Contribution of collagen and mineral to the elastic-plastic properties of bone. J Bone Joint Surg Am, 1975. 57(7): p. 956–61.

7. Hernandez, C.J. and T.M. Keaveny, A biomechanical perspective on bone quality. Bone, 2006. 39(6): p. 1173–81.

8. Oftadeh, R., et al., Biomechanics and mechanobiology of trabecular bone: a review. J Biomech Eng, 2015. 137(1): p. 0108021–01080215.

9. Orkoula, M.G., M.Z. Vardaki, and C.G. Kontoyannis, Study of bone matrix changes induced by osteoporosis in rat tibia using Raman spectroscopy. Vibrational Spectroscopy, 2012. 63: p. 404–408.

10. Monzem, S., et al., Raman spectroscopic of osteoporosis model in mouse tibia in vivo. Vibrational Spectroscopy, 2018. 98: p. 88–91.

11. Schulmerich, M.V., et al., Transcutaneous Raman Spectroscopy of Murine Bone In Vivo. Applied Spectroscopy, 2009. 63(3): p. 286–295.

12. Shu, C., et al., Spatially offset Raman spectroscopy for in vivo bone strength prediction. Biomedical Optics Express, 2018. 9(10): p. 4781–4791.

13. Massie, C., et al., Improved prediction of femoral fracture toughness in mice by combining standard medical imaging with Raman spectroscopy. J Biomech, 2021. 116: p. 110243.

14. Pejović-Milić, A., et al., Ultrasound measurements of overlying soft tissue thickness at four skeletal sites suitable for in vivo x-ray fluorescence. Med Phys, 2002. 29(11): p. 2687–91.

15. Meema, H.E. and H. Meindok, Advantages of peripheral radiogrametry over dual-photon absorptiometry of the spine in the assessment of prevalence of osteoporotic vertebral fractures in women. J Bone Miner Res, 1992. 7(8): p. 897–903.

16. Kiel, D.P., et al., Can metacarpal cortical area predict the occurrence of hip fracture in women and men over 3 decades of follow-up? Results from the Framingham Osteoporosis Study. J Bone Miner Res, 2001. 16(12): p. 2260–6.

17. Schreiber, J.J., R.N. Kamal, and J. Yao, Simple Assessment of Global Bone Density and Osteoporosis Screening Using Standard Radiographs of the Hand. J Hand Surg Am, 2017. 42(4): p. 244–249.

18. Buckley, K., et al., Towards the in vivo prediction of fragility fractures with Raman spectroscopy. Journal of Raman Spectroscopy, 2015. 46(7): p. 610–618.

19. Esmonde-White, F. and M. Morris, Validating in vivo Raman spectroscopy of bone in human subjects. SPIE BiOS. Vol. 8565. 2013: SPIE.

20. Maher, J.R. and A.J. Berger, Determination of Ideal Offset for Spatially Offset Raman Spectroscopy. Applied Spectroscopy, 2010. 64(1): p. 61–65.

21. Esmonde-White, F.W., K.A. Esmonde-White, and M.D. Morris, Minor distortions with major consequences: correcting distortions in imaging spectrographs. Applied spectroscopy, 2011. 65(1): p. 85–98.

22. Savitzky, A. and M.J.E. Golay, Smoothing and Differentiation of Data by Simplified Least Squares Procedures. Analytical Chemistry, 1964. 36(8): p. 1627–1639.

23. Maher, J.R., et al., Overconstrained library-based fitting method reveals age- and disease-related differences in transcutaneous Raman spectra of murine bones. Journal of Biomedical Optics, 2013. 18(7).

24. Chen, K., C. Massie, and A.J. Berger, Soft-tissue spectral subtraction improves transcutaneous Raman estimates of murine bone strength in vivo. Journal of Biophotonics, 2020. 13(n/a): p. e202000256.

